# L-type calcium channel blockade with verapamil prevents noise induced neuronal dys-synchrony

**DOI:** 10.1101/2024.07.10.602276

**Authors:** Selin Yalcinoglu, Rod D. Braun, Ammaar Wattoo, Aaron K. Apawu, Rasheed Alrayashi, Avril Genene Holt

## Abstract

Previous studies have established the protective effects of calcium channel blockade on the peripheral auditory system in response to noise exposure. While these studies implicate L-type calcium channels (LTCCs) in noise generated dysfunction in the auditory periphery, contributions of LTCCs to noise-induced central dysfunction remains unclear. To begin to elucidate the roles of LTCCs in hearing, peripheral and central auditory function were assessed longitudinally after LTCC blockade. Neuronal synchrony and activity were assessed by analyzing wave I (peripheral) and wave V (central) auditory brainstem responses (ABRs). Just prior to a noise exposure resulting in a temporary shift in hearing thresholds, rats were administered verapamil (LTCC blocker) or saline. Verapamil administration prevented the noise-induced decrease in ABR wave I and V amplitudes. Interestingly, when non-noise exposed animals were administered verapamil, wave V amplitude decreased, suggesting that LTCCs are critical for neuronal synchrony in the inferior colliculus. The inferior colliculus mediates inhibition of the acoustic startle reflex (giASR). Following noise exposure giASR was enhanced, but the enhancement was not prevented by LTCC blockade. These results suggest that while LTCCs are necessary for auditory-related synchronous activity, these channels do not contribute to noise-induced hyperactivity in the inferior colliculus.

---

Hearing loss affects approximately 48 million Americans, with more than half having lasting damage due to excessive noise exposure (Mitchell, 2006). Hearing loss can impact quality of life, cognition, and mental health with increased suicidal ideations (Blazer et al., 2016). Additionally, impaired communication with others can result in social isolation with fewer educational and job opportunities. These limitations lead to an estimated $3,000 in societal associated costs per affected person annually (Mohr et al., 2000; Stucky et al., 2010). Individuals whose occupational and recreational activities involve noisy environments are at higher risk for permanent or temporary hearing impairment (e.g., factory workers, military personnel, dental hygienists, musicians, or athletes) (Helzner et al., 2005; Mitchell, 2006). Regardless of the severity of the noise-induced hearing loss (NIHL), there can be sustained changes in peripheral and central auditory related function. Additionally, exposure to loud noise has been associated with synaptopathy, a loss of synaptic connection between the auditory nerve and inner hair cells in the periphery, and increased neuronal activity in central auditory brain regions, such as the inferior colliculus (Fernandez et al., 2015; Kaltenbach & Zhang, 2007; Kujawa & Liberman, 2009; Mulders & Robertson, 2009). Neuronal activity is primarily regulated by free calcium entering the neuron via voltage gated calcium channels and causing release of neurotransmitter (Clarke et al., 2016; Dolphin & Lee, 2020; Fukaya et al., 2023; Scarnati et al., 2020). The effects of increased calcium accumulation as a result of excessive neuronal activity can range from ototoxicity and cell death to increased spontaneous neuronal activity and have been linked to multiple auditory related disorders (Alharazneh et al., 2011; Orrenius et al., 2003; Szydlowska & Tymianski, 2010). Thus, regulation of calcium channel function might be a potential therapeutic approach to protect from both peripheral and central contributions to NIHL. Previous studies in rodents have used a variety of calcium channel antagonists to reduce the impact of a noise-induced permanent threshold shift in the periphery with some success (Bobbin et al., 1990a; Goldwyn & Quirk, 1997; Liu et al., 2012; Uemaetomari et al., 2009). However, there is still a gap in knowledge regarding the effects of calcium channel blockade on noise-induced synaptic function centrally. Verapamil, a non-dihydropyridine L-type calcium channel antagonist, has few side effects and has been shown to readily cross the blood-brain barrier to target calcium channels centrally (Colbourne & Harrison, 2022; Tfelt-Hansen & Tfelt-Hansen, 2009). Since, 1) voltage gated calcium channel function and calcium mobilization are crucial for normal auditory transmission (Kennedy, 2012; Liu et al., 2021; Purcell et al., 2011; Pyott et al., 2004; Zanazzi & Matthews, 2009, Zuccotti et al., 2013), and 2) noise exposure disrupts spontaneous neuronal activity (Bao et al., 2003; Cacace et al., 2014; Coomber et al., 2014; Dehmel et al., 2012; Hesse et al., 2016; Holt et al., 2005; Holt et al., 2010; Li et al., 2013; Muca et al., 2018; Niu et al., 2013), we hypothesize that blockade of these channels modulates neuronal synchrony and therefore provides protection from noise-induced trauma for both peripheral and central auditory transmission.

## Materials and Methods

### Subjects and design

Twenty-five young adult male Sprague-Dawley rats (Charles River Laboratory, New Jersey) were randomly divided into four treatments groups. The treatment groups were no noise exposure with saline (n=6), noise exposure with saline (n=7), no noise exposure with verapamil (n=5), and noise exposure with verapamil (n=7). Approximately 20 minutes prior to the noise exposure either verapamil (30 mg/kg) or saline (27 ml/kg, 0.9%) solution was administered intraperitoneally. The noise groups were exposed to a 16 kHz, 106 dB SPL tone for one hour, while no noise control groups were maintained in ambient noise conditions for an equal amount of time.

All rats were individually housed and maintained at 24 ± 1° C with a 12-h light/dark cycle (lights on at 7 AM). Standard housing conditions with free access to normal rat chow and tap water were provided at Wayne State University’s (WSU) AAALAC-accredited animal facility.

Animals were treated in accordance with the Animal Welfare Act and DHHS “Guide for the Care and Use of Laboratory Animals”, and all procedures were approved by the WSU Institutional Animal Care and Use Committee in accordance with NIH guidelines.

### Noise Exposure

Just prior to noise exposure a single ear of each animal was protected by filling the ear canal with a silicone elastomer (Kwik Seal, World Precision Instruments, Sarasota, FL). Awake freely moving animals were exposed to a 16 kHz tone, 106 dB SPL with a sound pressure level (SPL) of 106 dB generated using software (Daqarta v 10.3) and delivered by overhead speakers for one hour. The silicone plug was removed immediately following the noise exposure. Animals in the no noise group had one ear plugged for one hour but were not exposed to loud noise.

### Auditory Brainstem Responses (ABRs)

ABR analysis evaluated both the amplitudes (waves I and V) and hearing thresholds. To record the neurological response subdermal electrodes were placed below the test ear (reference), below the contralateral ear (ground), and at the vertex (active). The assessment was performed at two different frequencies (12 kHz and 20 kHz) at incremental intensities (between 30-85 dB) at three time points (same day, one day, and five days after treatment; Fig. 1). At each intensity amplitudes (μV) and latencies (ms) were calculated. Amplitudes were calculated by subtracting the negative peak from the corresponding positive peak of the response, and latencies were recorded at only the positive peak for each wave (I and V). Hearing thresholds were measured at 12 and 20 kHz, with decreasing increments beginning at 80 dB SPL, until a response was no longer elicited. The lowest intensity that elicited a response was recorded as the threshold.

**Figure 1.**
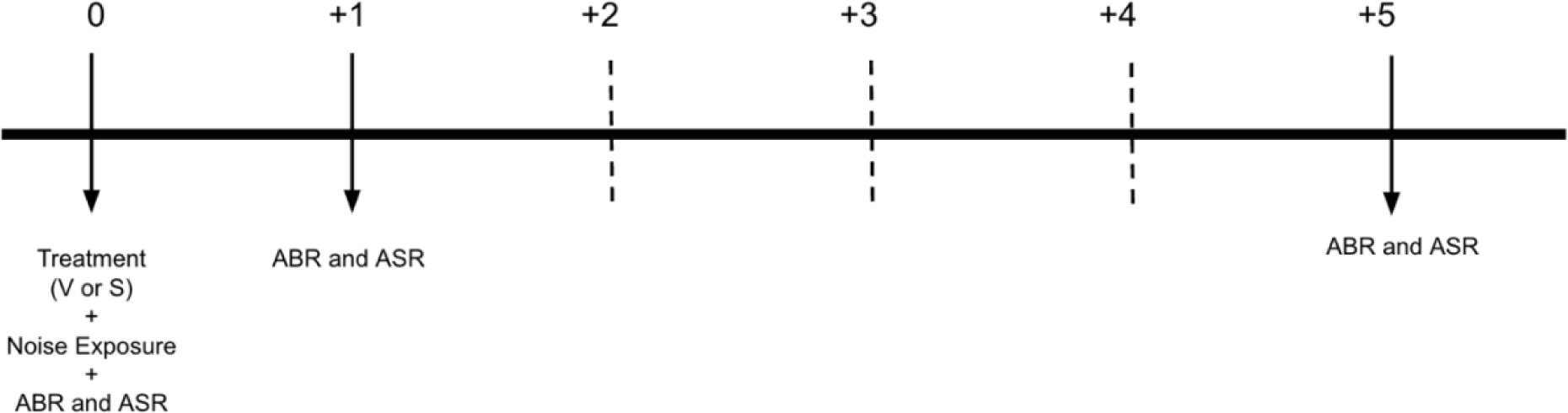
Experimental timeline. Following treatment (verapamil, V or saline, S) and noise exposure, each rat had ABR, and ASR tests performed at three different time points: same day after treatment (day 0), one day after treatment (+1 day), and five days after treatment (+5 days).

### Gap inhibition of Acoustic Startle Reflex (giASR)

giASR testing was performed at six frequencies (4, 8, 12, 16, 20 and 24 kHz) and two intensities (45- and 60-dB SPL). Testing was performed at three time points (same day, one day, and five days after treatment, Fig. 1). The gap inhibition of ASR paradigm was composed of three types of trials (Fig. 2): a silent period followed by a 20 ms startle sound (120 dB broadband noise; Fig. 2A, “sound-off startle” or “SOS” trial), a background tone immediately followed by a startle sound (Fig. 2B, “no gap” or “NG” trial), and a background tone interrupted by 50 ms of silence before reintroduction of the tone for 50 ms before the startle sound (Fig. 2C, “gap” or “G” trial). In each trial F_max_ (Newton), the maximum force exerted during the startle response, was recorded. Following the testing the F_max_ in gap trial to F_max_ in sound-off startle trial (G/SOS) ratio was analyzed.

**Figure 2.**
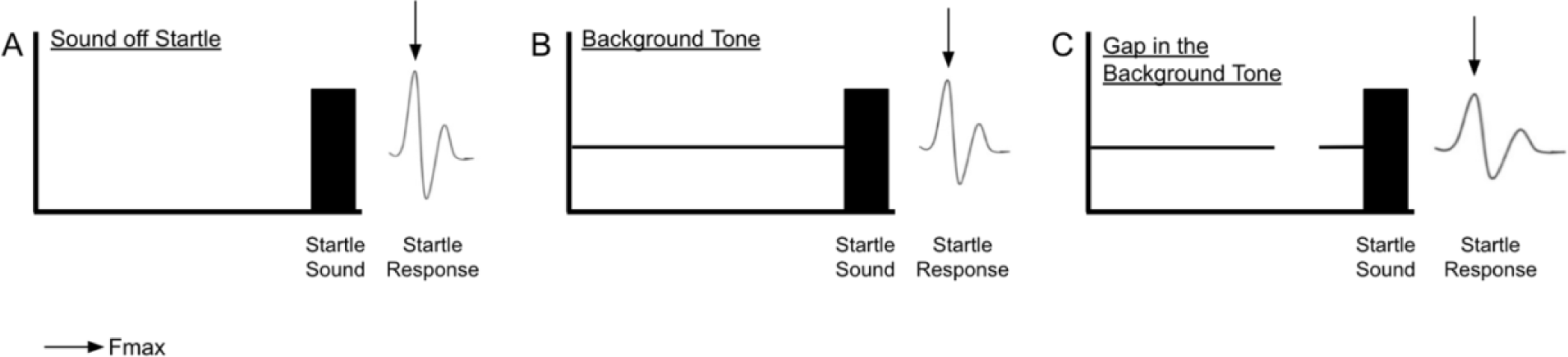
ASR testing paradigm. Gap detection was composed of three types of trials: a silent period followed by a startle sound (solid black bar), i.e., “sound-off startle” or “SOS” trial (A), a background tone immediately followed by a startle sound, i.e., “no gap” or “NG” trial (B), and a background tone interrupted by a 50 ms silent period before reintroduction of the tone for 50 ms before the startle sound, i.e., “gap” or “G” trial (C). In each trial F_max_, the maximum force exerted during the startle response, was recorded (indicated by the arrow).

### Statistical analysis

All data were analyzed using StatView Version 5.0.1.0 (SAS Institute Inc.) and GraphPad Prism version 10.0.0 for MAC (Boston, MA USA). Group differences were analyzed by analysis of variance (ANOVA) and Tukey/Kramer post-hoc comparison. P values less than 0.05 were considered significant. All data are expressed as mean ± standard error of the mean.

## Results

### ABR Analysis - Verapamil does not impact hearing thresholds in normal or noise exposed animals

#### Same day of treatment

On the same day of treatment (Fig. 3A), a significant threshold shift was observed in the noise saline group compared to no noise saline and no noise verapamil groups at 12 kHz (26 dB SPL and 31 dB SPL, respectively, p<0.05) and at 20 kHz (29 dB SPL, p<0.05). The noise verapamil group had a significant threshold shift compared to no noise saline and no noise verapamil at 12 kHz (28 dB SPL and 30 dB SPL, respectively, p<0.05) and at 20 kHz (33 dB SPL, p<0.05).

**Figure 3.**
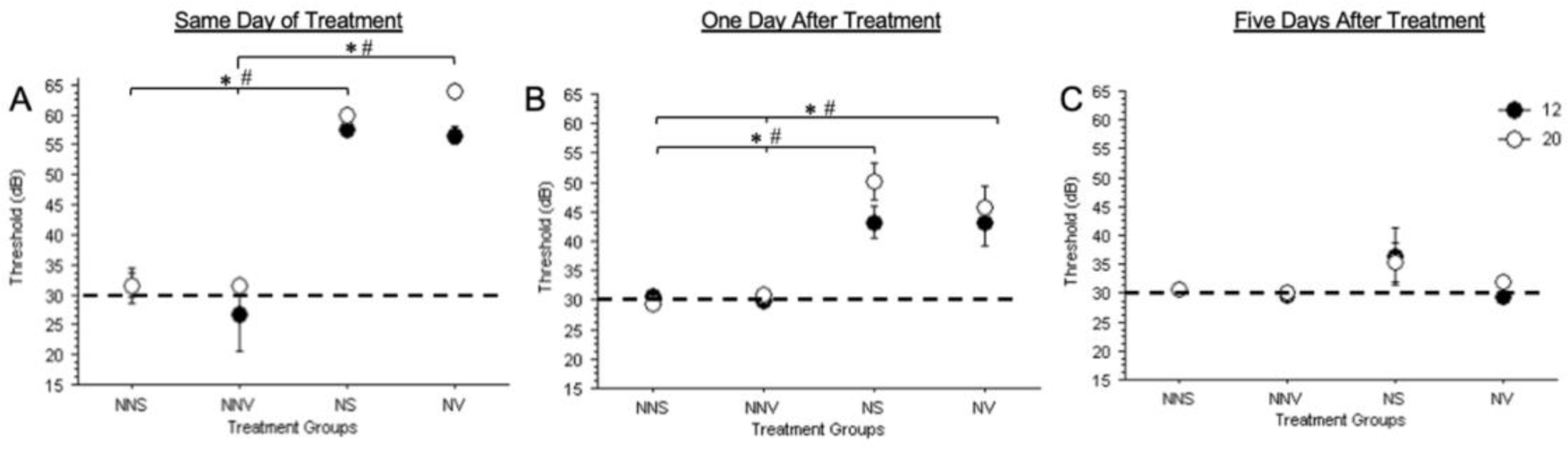
ABR thresholds at 12 and 20 kHz. Three different time points (A) same day of treatment, (B) one day after treatment, and (C) five days after treatment, are shown. Hearing thresholds of each treatment group (no noise saline, no noise verapamil, noise saline and noise verapamil) are displayed at 12 kHz (closed circle) and 20 kHz (open circle). Baseline hearing thresholds in normal hearing rats after either saline or verapamil administration are 30 dB SPL (indicated by the dashed line). Groups were compared using ANOVA with a post-hoc Tukey-Kramer test, and the error bars indicate the standard error of mean. * Asterisks denote significance at 12 kHz and # hashtag denote significance at 20 kHz at p<0.05.

#### One day after treatment

One day after treatment, thresholds were still elevated in noise groups both at 12 and 20 kHz (Fig. 3B). A significant threshold shift was observed in the noise saline group compared to no noise saline and no noise verapamil groups at 12 kHz (13 dB SPL, p<0.05) and at 20 kHz (21 dB SPL and 19 dB SPL, respectively, p<0.05). The noise verapamil group had a significant threshold shift compared to no noise saline and no noise verapamil at 12 kHz (13 dB SPL, p<0.05) and at 20 kHz (16 dB SPL and 15 dB SPL, respectively, p<0.05).

#### Five days after treatment

Five days after noise exposure (Fig. 4C) hearing thresholds of both the noise groups returned to baseline with no significant difference compared to no noise groups.

**Figure 4.**
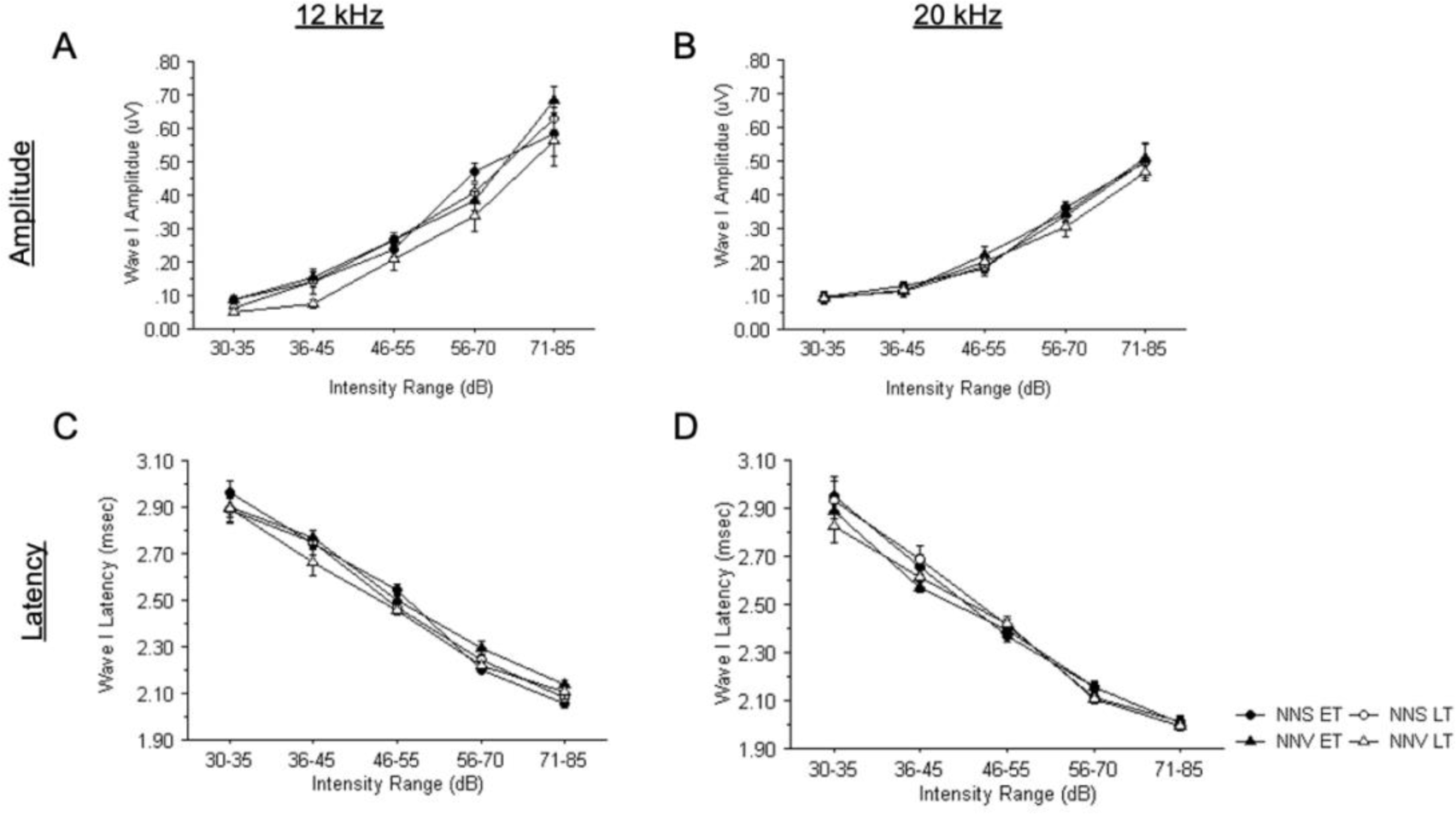
ABR wave I amplitude and latency in relation to intensity at 12 and 20 kHz. Wave I amplitude (A-B) and latency (C-D) across 5 intensity ranges are displayed. The wave I amplitudes and latencies were analyzed for no noise saline early time points (closed circles), no noise saline late time points (open circles), no noise verapamil early time points (closed triangles), no noise verapamil late time points (open triangles), groups at 12 kHz (A, C) and 20 kHz (B,D).

### Verapamil does not interrupt expected wave I input / output Function

#### Amplitude

At early time points (days 0 and 1) wave I amplitude at both 12 and 20 kHz (Fig. 4A and 4B respectively) increases as intensity of the tone increases. At the late time point (day 5) wave I amplitude increases as intensity of the tone increases at both 12 and 20 kHz (Fig. 4A and 4B respectively)

#### Latency

At early time points (days 0 and 1) wave I latency at both 12 and 20 kHz (Fig. 4C and 4D respectively) decreases as intensity of the tone increases. At the late time point (day 5) wave I latency decreases as intensity of the tone increases at both 12 and 20 kHz (Fig. 4C and 4D respectively).

### Verapamil prevents noise induced synaptopathy at suprathreshold

#### 12 kHz

On day zero of noise exposure there was a significant wave I amplitude reduction in the noise saline group compared to the no noise saline and the no noise verapamil groups (84% and 86% respectively, p<0.05) at 12 kHz (Fig. 5A). The noise verapamil group was not significantly different when compared to the no noise saline and the no noise verapamil groups. One day after the noise exposure wave I amplitudes returned to normal for the noise saline group with no significant difference between the no noise groups (Fig. 5A). In addition, verapamil did not have any impact on wave I amplitude at later time points (Fig. 5A).

**Figure 5.**
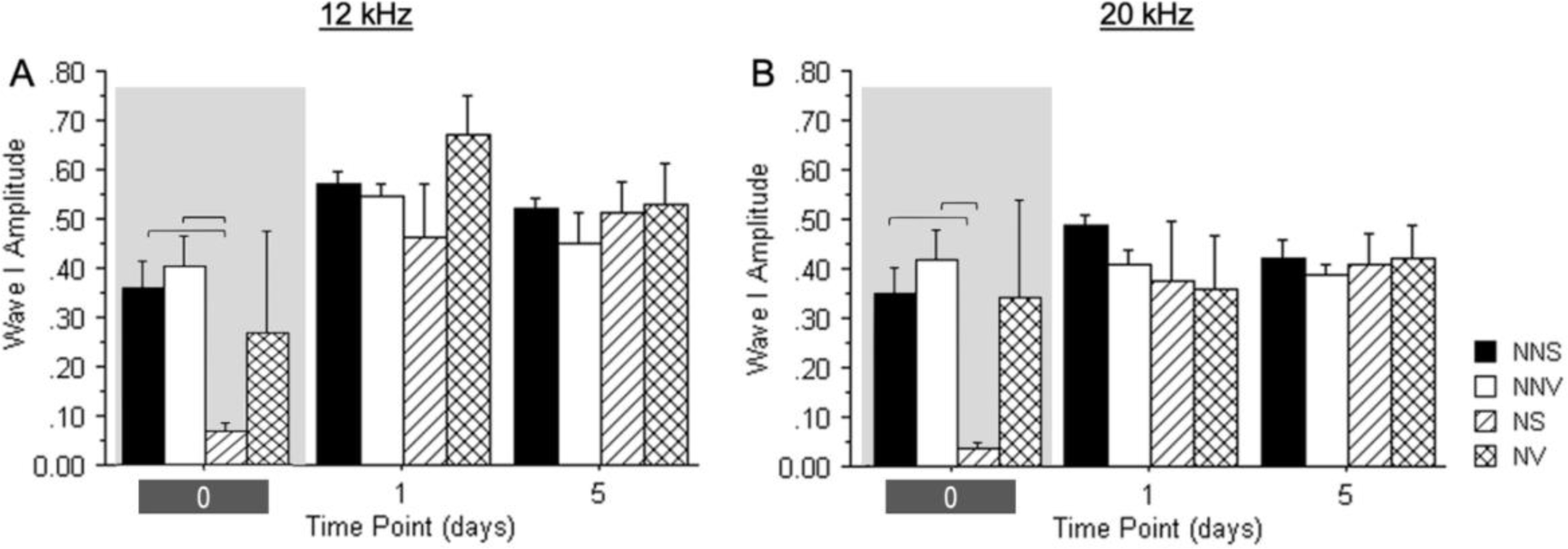
ABR wave I amplitude for auditory brainstem responses at 12 and 20 kHz. Wave I amplitudes on day zero, and one and five days after treatment and noise exposure are displayed. The wave I amplitudes were analyzed for no noise saline (black solid bars), no noise verapamil (solid white bars), noise saline (hatched bars) and noise verapamil (crossed bars) groups at 12 kHz (A) and 20 kHz (B). Groups were compared using ANOVA with a post-hoc Tukey-Kramer test, and the error bars indicate the standard error of mean. Brackets denote significance at p<0.05. Shaded set of bars indicate the time point at which significant changes were observed.

#### 20 kHz

On day zero of noise exposure there was a significant wave I amplitude reduction in the noise saline group compared to the no noise saline and the no noise verapamil groups (77% and 79% respectively, p<0.05) at 20 kHz (Fig. 5B). The noise verapamil group was not significantly different when compared to the no noise saline and the no noise verapamil groups (Fig. 5B). One day after the noise exposure wave I amplitudes returned to normal for the noise saline group with no significant difference between the no noise groups (Fig. 5B). In addition, verapamil did not have any impact on wave I amplitude at later time points (Fig. 5B).

### Verapamil does not interrupt expected wave V input / output Function

#### Amplitude

At early time points (days 0 and 1) wave V amplitude at both 12 and 20 kHz (Fig. 6A and 6B respectively) increases as intensity of the tone increases. At the late time point (day 5) wave I amplitude increases as intensity of the tone increases at both 12 and 20 kHz (Fig. 6A and 6B respectively)

**Figure 6.**
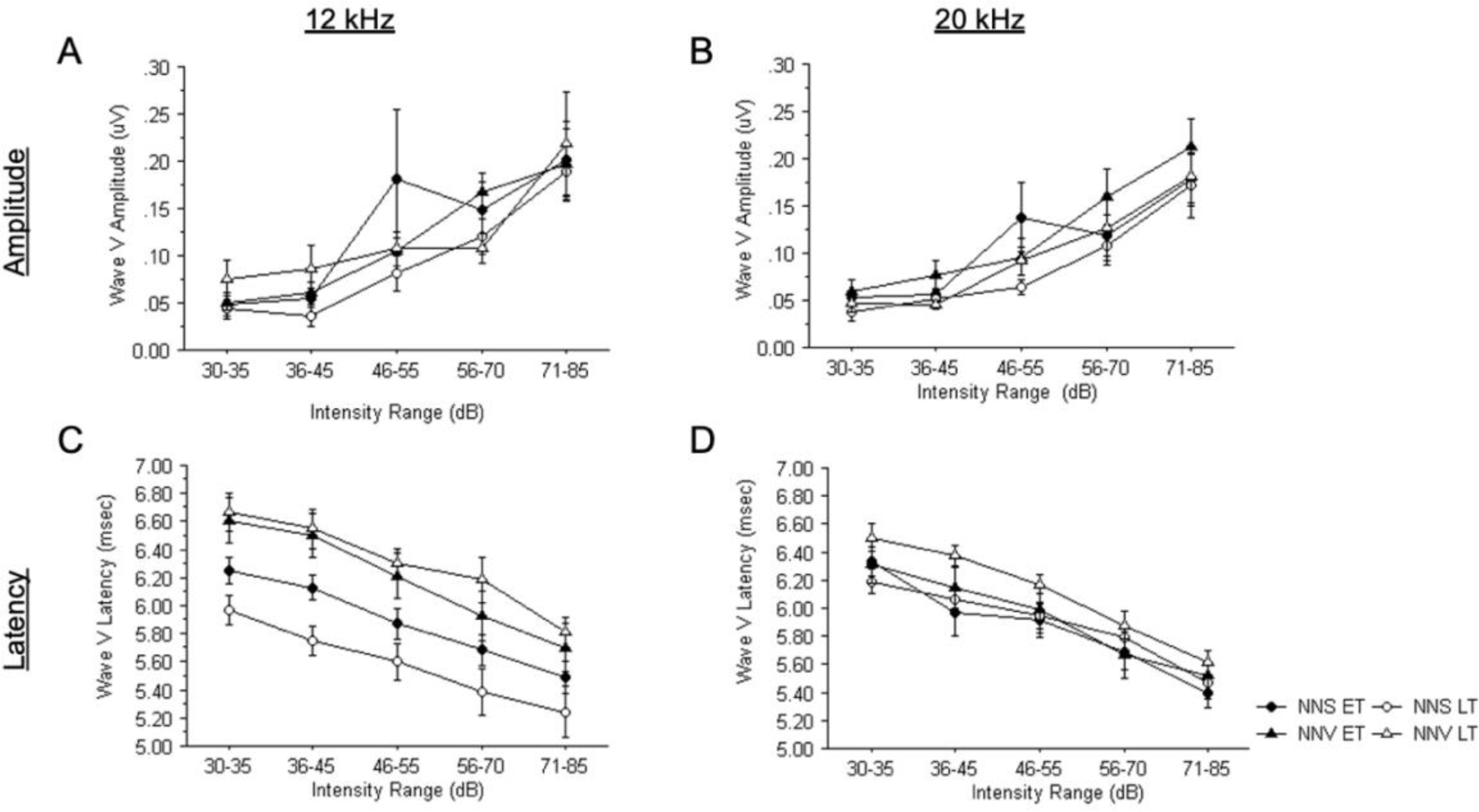
ABR Wave V amplitude and latency in relation to intensity at 12 and 20 kHz. Wave V amplitude (A-B) and latency (C-D) across 5 intensity ranges are displayed. The wave V amplitudes and latencies were analyzed for no noise saline early time points (closed circles), no noise saline late time points (open circles), no noise verapamil early time points (closed triangles), no noise verapamil late time points (open triangles), groups at 12 kHz (A, C) and 20 kHz (B, D).

#### Latency

At early time point (days 0 and 1) wave V latency at both 12 and 20 kHz (Fig. 6C and 6D respectively) decreases as intensity of the tone increases. At the late time point (day 5) wave I latency decreases as intensity of the tone increases at both 12 and 20 kHz (Fig. 6C and 6D respectively)

### The impact of Verapamil on central synchronous activity is dependent on noise exposure status

#### 12 kHz

On day zero of noise exposure there is a trend towards wave V amplitude reduction in the noise saline group compared to no noise saline group at 12 kHz (Fig. 5A). In addition, no noise verapamil group had a non-detectable wave V amplitude on day zero of treatment. However, on day zero of noise exposure, there is no wave V amplitude reduction in the noise verapamil group compared to the noise saline group. One day after the noise exposure and treatment the reduction observed in the no noise verapamil group on day zero significantly increased (p<0.001) and was not significantly different compared to other groups.

#### 20 kHz

On day zero of noise exposure there is a trend towards wave V amplitude reduction in the noise saline group compared to the no noise saline group at 20 kHz (Fig. 5B). However, on day zero of noise exposure a similar trend of a reduction in wave V amplitude in the noise verapamil group compared to the no noise saline group was not observed. One day after the noise exposure wave V amplitudes returned to normal for all noise groups with no significant difference between the no noise groups.

### Verapamil enhances noise induced inhibition of the acoustic startle reflex

#### G/SOS ASR Ratio at 45 dB

On day zero of treatment, the noise verapamil group showed a decrease in G/SOS ratio compared to the no noise saline (46% 12 kHz p<0.05) and no noise verapamil (38%, at 12 kHz p<0.05) group on day zero of treatment (Fig. 9A). By one day after treatment, the noise verapamil group showed a decrease in G/SOS ratio compared to the noise saline group at 20 kHz (47%, p<0.05, Fig. 9B). Five days after treatment noise verapamil group showed an increase in G/SOS ratio compared to no noise verapamil group at 20 kHz (33%, p<0.05, Fig. 9B).

#### G/SOS ASR Ratio at 60 dB

On day zero of treatment, the noise saline group showed a decrease in G/SOS ratio compared to the no noise saline group (65% and 54% at 12 and 20 kHz respectively p<0.05 Fig. 9 C-D) and no noise verapamil (50% at 20 p<0.05, Fig. 9 D). The noise verapamil group showed a decrease in G/SOS ratio compared to the no noise saline and no noise verapamil (70%, and 63% respectively, p<0.05) groups on day zero of treatment at 12 kHz (Fig. 9 C). By one day after treatment the noise verapamil group had significantly lower G/SOS ratios compared to no noise verapamil (63%, at 20 kHz, respectively, p<0.05, Fig. 9 D). Five days after treatment noise verapamil group showed deficits in G/SOS ratio compared to no noise saline group at 20 kHz (19%, p<0.05, Fig. 9D).

## Discussion

The current study demonstrates the protective effects of L-type calcium channel blockade on neuronal synchrony in a model of temporary hearing loss. L-type calcium channel blockade prior to noise exposure prevented peripheral and central disruptions in neuronal synchrony, demonstrated by decreases in ABR wave I and V amplitudes respectively. The results also showed that L-type calcium channel blockade, independent of noise overexposure, reduced neuronal synchrony in midbrain regions such as the inferior colliculus. However, even deficits in midbrain neuronal synchrony did not disrupt inferior colliculus mediated inhibition of the acoustic startle reflex (giASR). Conversely, normal synchronous activity in the periphery does not prevent pathological giASR performance. This demonstrates that giASR performance is not dependent on ABR detectable peripheral or central neuronal synchrony.

### L-type Voltage gated calcium channel blockade does not prevent noise induced temporary threshold shift

Previous studies have shown that calcium channels play an integral role in cochlear function (Bobbin et al., 1990b; Liu et al., 2012; Liu et al., 2021). Protective effects of L-type and T-type calcium channel blockade have been shown in response to a noise exposure that typically produces a permanent hearing loss. In the current study we explored the impact of L-type calcium channel blockade in a model of temporary hearing loss, but unlike the permanent threshold model, in this milder noise exposure model, hearing loss was not prevented. The mechanism underlying temporary versus permanent hearing loss is still unclear, however the findings from the current study suggest that L-type calcium channels are not primary contributors to noise induced temporary hearing loss. Other studies have implicated loss of tip links (Hazra et al., 2019; Kazmierczak et al., 2007; Le et al., 2017; Mao & Chen, 2021; Sabouni Tabari et al., 2022; Siemens et al., 2004; Wang et al., 2018). Although calcium channel blockade does not significantly affect hearing thresholds, noise exposure may produce synaptopathy (diminished synchronous activity) which may be present despite recovery of the hearing thresholds.

### Calcium channel blockade prevents noise induced neuronal dys-synchrony peripherally

Auditory function can be assessed throughout the auditory pathway by analyzing individual peaks present in the ABR wave form (Bachmann et al., 2024; Chang et al., 2022; Jiang et al., 2021; Wood et al., 2019). For example, peripheral auditory function corresponds to the first peak (Wave I) of the ABR, present at about 2 msec from the onset of the stimulus (Chang et al., 2022; Wood et al., 2019). Disruption to the neural connections (synapses) between the inner hair cells (IHC) and the auditory nerve has been associated with changes in Wave I of the ABR. Cochlear synaptopathy, loss of synaptic connections with IHCs or auditory nerve: IHC synaptic dysfunction, may manifest as a decrease in ABR Wave I amplitude following noise trauma (Bramhall et al., 2017; Greguske et al., 2021; Hashimoto et al., 2019; Ingersoll et al., 2020; Kujawa & Liberman, 2015; McFarlane & Sanchez, 2023; Prendergast et al., 2017). Additionally, overstimulation of IHCs, as is the case during noise over exposure, can lead to increased calcium influx, which if not cleared from the ribbon can be toxic, leading to synaptic swelling and bursting, and possibly cell death (Altschuler et al., 2016; Beurg et al., 2008; Grant & Fuchs, 2008; Liang et al., 2019; Liu et al., 2021; Maurer et al., 1999; Pyott et al., 2004; Zachary et al., 2018; Zhang et al., 2018). This loss of synaptic connections or disruption in synchronous firing of neurons in response to the stimulus following a loud noise exposure may result in decreases in ABR Wave I amplitude.

A single loud noise exposure can lead to hearing loss at frequencies different from that of the noise exposure frequency (Bakay et al., 2018; Carroll et al., 2017; Choi et al., 2012; Daniel, 2007; Hong et al., 2015). Therefore, when assessing the impact of a single noise exposure, frequencies both above and below the noise exposure frequency were analyzed. Though synaptic counts were not performed in the current study, the reduction observed in ABR Wave I amplitude following noise exposure suggests that there is a loss of neural synchrony in the periphery at both 12 kHz and 20 kHz (Fig. 5) given that the presence of ABR waveforms is dependent on synchronous neuronal firing. Similar to other neuronal pathways in the nervous system, synaptic activity in the peripheral auditory system is regulated by calcium (Kennedy, 2012; Liu et al., 2021; Purcell et al., 2011; Pyott et al., 2004; Zanazzi & Matthews, 2009), thus calcium levels and calcium channel function can impact neural synchrony. Therefore, predictably, calcium channel blockade prior to noise exposure retained neuronal synchrony at the auditory nerve: IHC synapse (Fig. 5). The protective effect of calcium channel blockade was observed at both low (Fig. 5A) and high (Fig. 5B) frequencies. Since verapamil is lipophilic and can cross the blood brain barrier, the effects of verapamil observed in the periphery may extend centrally.

### Calcium channel blockade prevents noise induced neuronal dys-synchrony centrally

Central auditory function has been correlated with peaks II-V of the ABR wave form. Specifically, the neuronal synchronous activity in the inferior colliculus can be assessed by analyzing peak V (Wave V) at approximately 6 msec from the onset of the stimulus (Bachmann et al., 2024; Chang et al., 2022; Jiang et al., 2021; Wood et al., 2019). Similar to the peripheral effects of noise over exposure, a loss of neuronal synchrony was observed centrally, as indicated by decreases in ABR Wave V amplitude at both 12 and 20 kHz. These reductions in ABR Wave V amplitude, however, may not reflect hypofunction. In previous studies, spontaneous neuronal activity in the auditory pathway can change following noise exposure depending on the time after the noise (Cacace et al., 2014; Holt et al., 2010; Muca et al., 2018) Noise induced increases in activity within the inferior colliculus have been demonstrated using manganese enhanced MRI (MEMRI) (Cacace et al., 2014; Holt et al., 2010; Muca et al., 2018) Since the paramagnetic manganese ion can be taken up by neurons via L-type voltage gated calcium channels, manganese uptake can identify brain regions with more active neurons, with MEMRI maps reflecting high and low regions of activity. Therefore, the fact that neuronal synchrony is disrupted, does not disagree with studies demonstrating increased spontaneous hyperactivity following noise exposure. Indeed, neurons with hyper normal activity, such as those found in the inferior colliculus following noise exposure, may not show the same dys-synchronous discharge patterns as before the noise trauma.

### Calcium channel blockade disrupts neuronal synchrony centrally

Interestingly, calcium channel blockade, in the absence of noise exposure, produces a loss of neuronal synchrony, as demonstrated by a significant decrement in ABR Wave V (Fig. 7-8). Since the neuronal synchrony in the periphery is not impacted by calcium channel blockade in the absence of noise exposure, this suggests that the inferior colliculus may be a region in the auditory pathway with high levels of LTCCs that are active at rest (Berger & Bartsch, 2014; Clarke & Lee, 2018; Drotos et al., 2024; Hanson & Hurley, 2016; Leaver et al., 2016; Slugocki et al., 2017). Therefore, synchronous activity would be disproportionately affected by calcium channel blockade in the inferior colliculus centrally, compared to the auditory periphery. While calcium channel blockade did not affect hearing thresholds, calcium channel blockade induced changes in central synaptic function can have effects on speech recognition and processing (Buschges et al., 2000; Choi et al., 2024; Cucis et al., 2021; Delaney & Crane, 2016; Ding & Vaziri, 1998; Friesen et al., 2001; Fu & Nogaki, 2005; Henry & Turner, 2003; Hildebrand et al., 2004; Ricoy & Frerking, 2014; Wheeler et al., 1994) This would suggest that use of calcium channel blockers for treatment of hypertension may generate undiagnosed synaptopathy.

**Figure 7.**
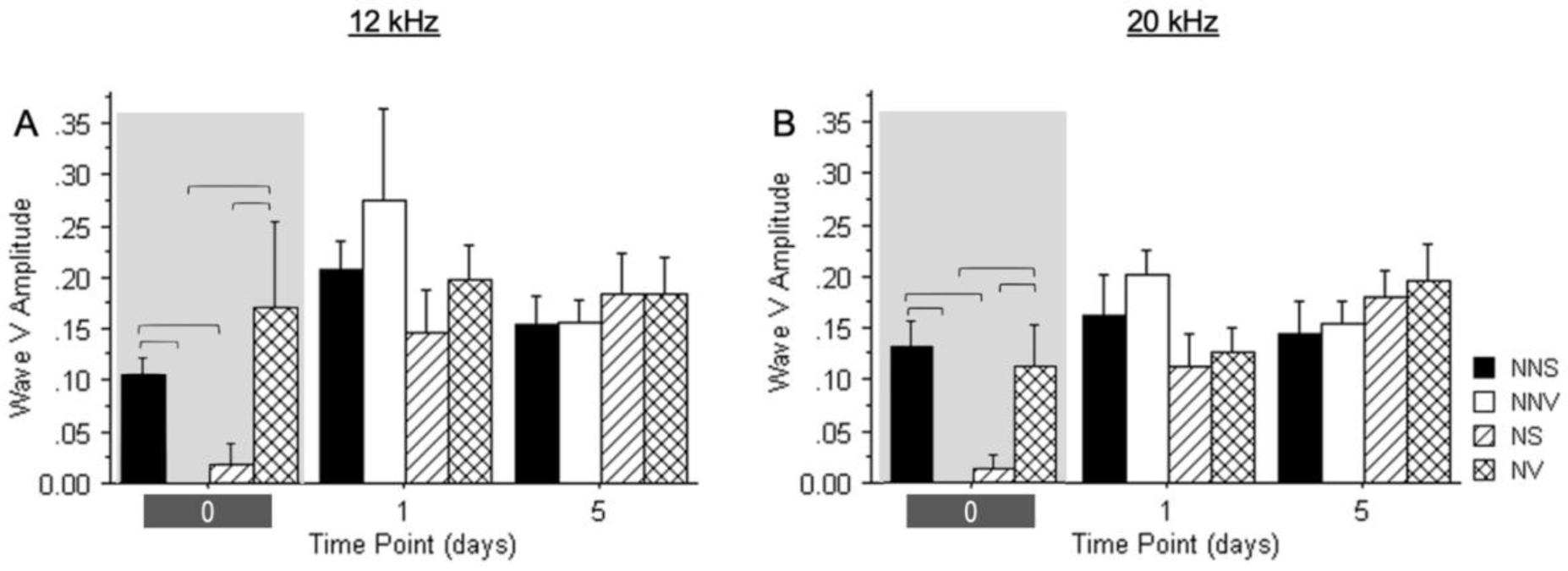
ABR wave V amplitude for auditory brainstem responses at 12 and 20 kHz. Wave V amplitudes on day zero-, one- and five-day post treatment and noise exposure are displayed. The wave V amplitudes were analyzed for no noise saline (black solid bars), no noise verapamil (solid white bars), noise saline (hatched bars) and noise verapamil (crossed bars) groups at 12 kHz (A) and 20 kHz (B). Groups were compared using ANOVA with a post-hoc Tukey-Kramer test, and the error bars indicate the standard error of mean. Brackets denote significance at p<0.05.

**Figure 8.**
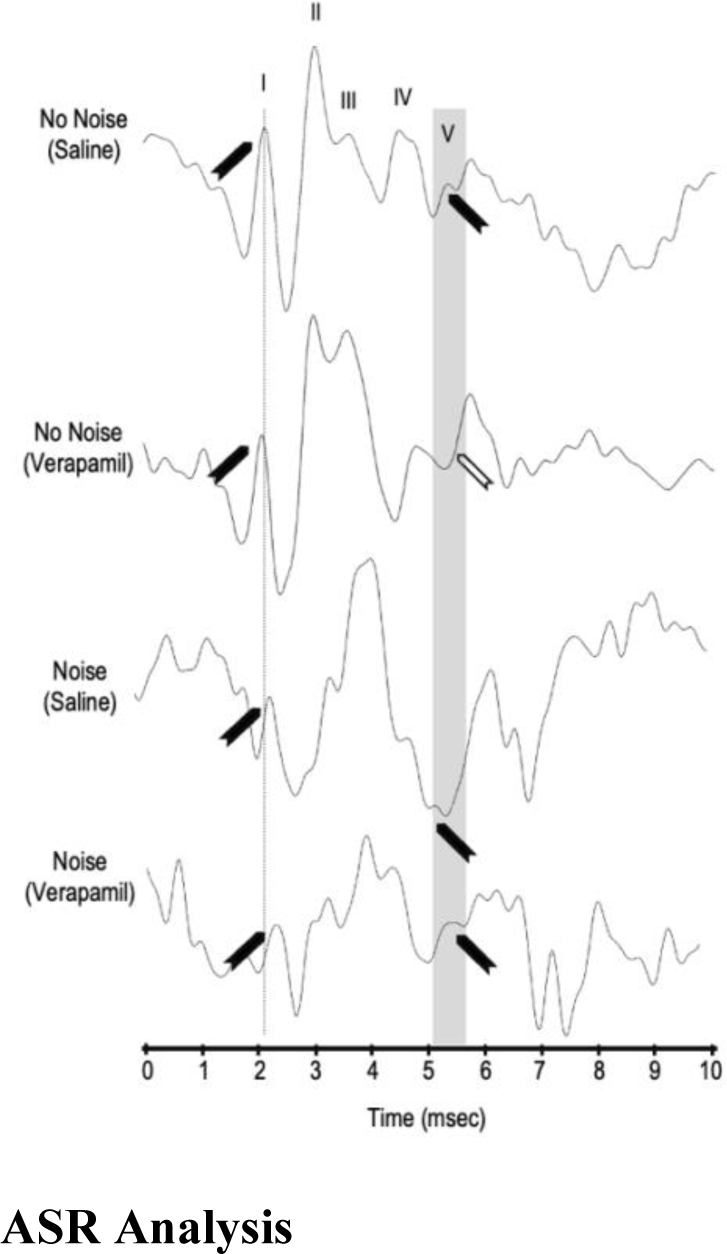
Auditory Brain Stem Response wave forms at 12 kHz at 80 dB. Response to a tone pip at day 0 of no noise (saline or verapamil), and noise (saline or verapamil) groups are displayed. Wave I, II, III, IV, V are indicated on the no noise saline group. Wave I is indicated by a dashed line and wave V is indicated by a gray bar. Waves that are present are marked by solid black arrowhead and waves that are absent are marked by a white arrowhead.

However, calcium channel blockade just prior to noise exposure, protected against noise-induced changes in neuronal synchrony. This may be due to noise induced changes in calcium channel density or localization. Another possibility is that reduced access to extra cellular calcium following calcium channel blockade triggers a secondary calcium mobilization pathway.

### Calcium channel blockade may activate secondary calcium mobilization pathways

The loss of synchronous activity observed centrally following calcium channel blockade or noise exposure is not sustained when calcium channel blockade is present at the time of noise exposure (Fig. 7). This may be due to the release of intracellular stores of calcium to compensate for decreased extracellular calcium mobilization. Ryanodine receptors (RyRs) are involved in the release of intracellular calcium from the endoplasmic reticulum (Ni et al., 2007; Santulli et al., 2017; Seo et al., 2015; Xie et al., 2019; Zissimopoulos et al., 2006)There are three isoforms of RyRs (RyR −1, −2, −3) with unique distributions in skeletal muscle, cardiac muscle, and brain (Jeyakumar et al., 1998; Wang et al., 2001; Zieminska et al., 2006). Each RyR isoform is expressed in the brain with RyR3 being most abundant (Adasme et al., 2011; Bruno et al., 2012; Bull et al., 2008; Casas-Tinto et al., 2011; Giannini et al., 1995; Meissner, 2017). Unlike skeletal ryanodine receptors, activation of cardiac ryanodine receptor (RyR-2) is solely dependent on intracellular levels of calcium and is independent of the membrane depolarization (Perni, 2022) Although what is known about the structure and function of RyR-3 is limited, biochemical and structural analyses have demonstrated the greatest similarities of this isoform with RyR-2 (Lanner et al., 2010; McPherson & Campbell, 1993; Van Petegem, 2012). Therefore, RyR-3 isoform may also be dependent on intracellular levels of calcium and may not interact with dihydropyridine receptors, such as L-Type voltage gated calcium channels. Intracellular mechanisms for calcium release may be the underlying process when extracellular calcium influx the cell is blocked, as in the current study. A change in the calcium mobilization in the inferior colliculus may lead to behavioral changes. Since the inferior colliculus plays a critical role in many acoustically evoked reflexes, changes to calcium trafficking can manifest as changes observed in these reflexes (Berger & Coomber, 2015) Therefore, the impact of calcium channel blockade on synchronous auditory synaptic function and calcium mobilization centrally, may lead to changes in giASRs.

### L-type calcium channel blockade does not prevent noise-enhanced giASR

To assess temporal auditory processing, giASRs are evaluated (i.e., the subconscious response of an awake behaving animal in response to a startling sound preceded by a silent gap embedded in a noise) (English & Drummond, 2021; Fournier & Hebert, 2016; Ison et al., 1991; Kodsi & Swerdlow, 1995; Lobarinas et al., 2013; Sebo et al., 2020; Shi et al., 2022; Sun et al., 2011; Weible et al., 2020).In line with previous studies, inhibition of the acoustic startle reflex (ASR) is observed in the presence of a silent gap (Fournier & Hebert, 2016, 2021; Galazyuk & Hebert, 2015; Longenecker et al., 2021; Moreno-Paublete et al., 2017; Weible et al., 2020; Wilson et al., 2019). This inhibition of the ASR is not diminished following noise exposure, calcium channel blockade or combination of both (Fig.10). In fact, an enhanced ability to inhibit the ASR is observed following noise exposure and calcium channel blockade prior to noise exposure (Fig. 10). This enhanced performance, despite disruption to synchronous neuronal activity observed with ABRs, suggests that giASR responses may not be dependent on neuronal synchrony from inner hair cell: auditory nerve synapses or inferior colliculus synapses. Additionally, when ABR derived synchronous neuronal activity recovered after noise exposure (Fig.10), giASR performance declined for noise exposed animals administered verapamil (Fig.10). The acoustic startle pathway (coronal sections modified from Paxinos and Watson 2007 6^th^ Ed.) involves the cochlear nucleus and the caudal pontine reticular nucleus (Fig.10). Activation of both nuclei increases motor neuron activity, resulting in the startle reflex. However, gap inhibition of acoustic startle reflex engages the inferior colliculus (Fig.10). The contribution of inferior colliculus to giASR pathway has been established, with lesioning of the inferior colliculus resulting in reduced inhibition of the acoustic startle reflex (e.g., increased startle (Campbell et al., 2007; Davis et al., 1982; Leaton & Brucato, 2001; Li et al., 1998; Lu et al., 2011; Lyall et al., 2009; Yeomans et al., 2010). When the inferior colliculus is engaged, there may be excitatory input to the pedunculopontine tegmental nucleus and a subsequent increase in the acetylcholine release to the caudal pontine reticular nucleus (Noftz et al., 2020; Schofield et al., 2011; Zhao et al., 2022). The reduced activity in the caudal pontine reticular nucleus leads to decreased motor neuron activity resulting in the inhibition of the acoustic startle reflex. In the current study, both the enhanced ASR inhibition following noise exposure and the subsequent reduced ASR inhibition five days after calcium channel blockade in the noise group may reflect changes in neuronal activity in the inferior colliculus.

**Figure 9.**
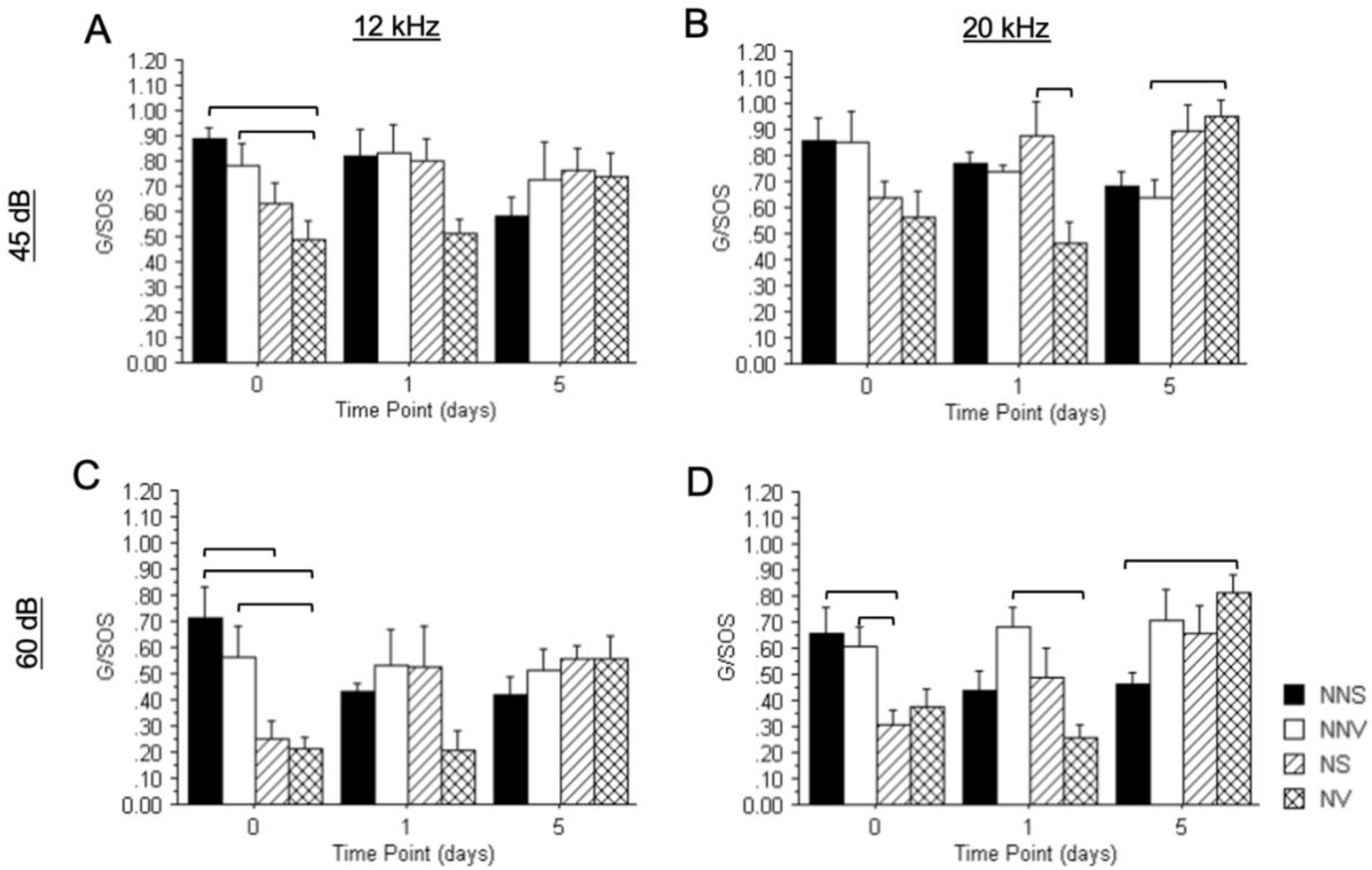
Gap to sound off startle ASR ratios at 12, and 20 at 45- and 60-dB SPL. The gap to sound off startle ASR ratios at three different time points, on day zero-, one- and five-day post treatment and noise exposure are displayed. The gap to sound off startle ASR ratios were analyzed for no noise saline (black solid bars), no noise verapamil (solid white bars), noise saline (hatched bars) and noise verapamil (crossed bars) groups at 12 kHz (A, C), and 20 kHz (B, D) at 45 dB (A, B) and 60 dB (C, D). Groups were compared using ANOVA with a post-hoc Tukey-Kramer test, and the error bars indicate the standard error of mean. Brackets denote significance at p<0.05.

**Figure 10.**
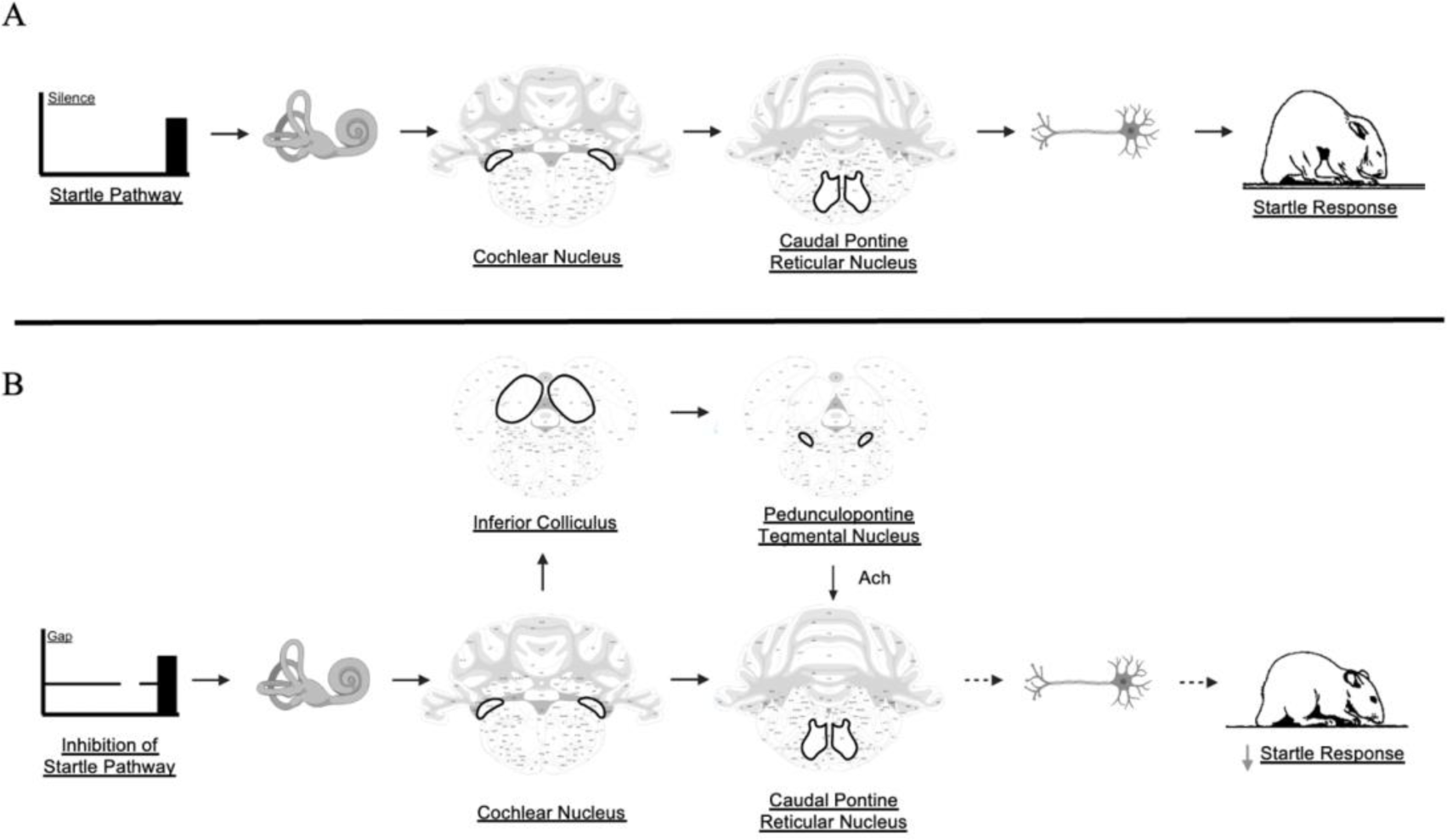
Acoustic startle reflex and gap inhibition of acoustic startle pathway. The startle reflex involves the activation of the caudal pontine reticular nucleus and subsequent activation of motor neurons (A). Gap inhibition of the acoustic startle reflex (giASR) engages the inferior colliculus (B). Increase of inferior colliculus activity increases the acetylcholine release from the pedunculopontine tegmental nucleus to the caudal pontine reticular nucleus and a subsequent decrease in motor neuron activity. This ultimately leads to inhibition of the acoustic startle reflex.

### Differential correlation of ABR waves with giASR performance

In the current study we assessed ABR Wave I and Wave V to determine whether amplitude correlated with giASR performance (Fig. 11). Reduced ABR Wave I amplitude observed following noise overexposure has been associated with poor ASR inhibition during giASR testing. These results correlate with other studies showing decreased Wave I amplitude after noise (Ruttiger et al., 2013; Bramhall et al., 2017; Bramhall et al., 2022; Mehraei et al., 2016; Nam et al., 2021; Prendergast et al., 2017; Rouse et al., 2020). In the current study, enhanced giASR performance has been observed after exposure to loud noise. For ABR Wave I, there was no significant correlation between amplitude and giASR ratios (i.e., G/SOS) in response to 12 or 20 kHz, 45 dB SPL carrier tones (R^2^= 0.0029 and 0.0801 respectively; data not shown). A similar result was found when the carrier tone was 12 or 20 kHz, 60 dB SPL (R^2^= 0.0094 and 0.1247 respectively, Fig. 11A-B). While there was no significant correlation between ABR Wave V amplitudes and giASRs elicited with 12 or 20 kHz, 45 dB carrier tones (R^2^= 0.2074 and 0.1916 respectively, data not shown), there was a significant correlation between ABR Wave V amplitudes and G/SOS in response to 12 or 20 kHz, 60 dB carrier tones (R^2^= 0.4114 and 0.2712 respectively, *p*≤0.05 Fig. 11 C-D). As the neuronal synchrony in ABR Wave V decreased, giASR performance was enhanced. This suggests that although the synchronous activity of the neurons is reduced following noise, overall levels of neuronal activity may increase. The inverse correlation between Wave V amplitude and enhanced giASR performance supports the premise that noise trauma produces dys-synchrony and hyperactivity in the inferior colliculus.

**Figure 11.**
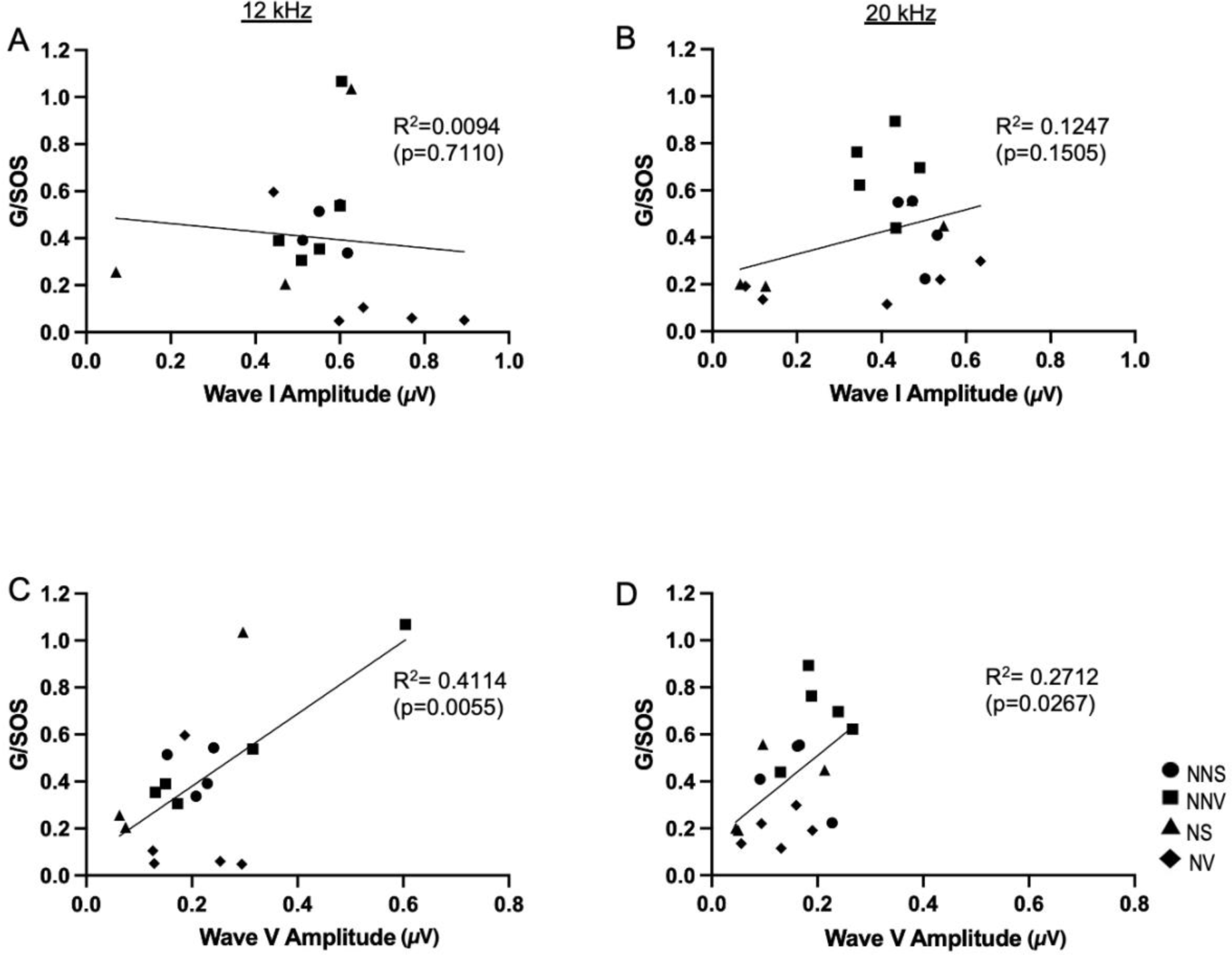
G/SOS – 60 dB SPL in relation to ABR Wave I and V amplitude at 12 and 20 kHz. Correlation of Wave I amplitude (A-B) and Wave V amplitude (C-D) and giASR (G/SOS) are displayed. The correlation was assessed at one day across groups no noise saline (circle - NNS), no noise verapamil (square - NNV), noise saline (triangle - NS) and noise verapamil (diamonds - NV). Significance of the slopes was determined using a regression t-test; p-values are shown.

### Conclusions

The present study demonstrates that voltage-gated calcium channels differentially contribute to peripheral and central auditory function. Blockade of LTCCs does not appear to disrupt neuronal activity (hearing thresholds and giASRs are unchanged). Additionally, this blockade results in a loss of neuronal synchrony centrally. Surprisingly, noise decreases neuronal synchrony both peripherally and centrally and verapamil prevents this outcome. Overall, these findings suggest calcium channel dysfunction may be an underlying mechanism for disorders involving hypo-or hyper-synchrony, but not neuronal hyperactivity. Calcium mobilization, whether by acoustic overexposure or calcium channel blockade may lead to an imbalance of extracellular and intracellular calcium ions. Therefore, exploring relationships between neuronal synchrony, neuronal hyperactivity, and calcium mobilization may enhance our future understanding of mechanisms and aid in identification of therapeutic approaches related to noise induced auditory dysfunction.

## Acknowledgments

We thank Mirabela Hali, Batoul Ayad, Peter Kamash, and Nyla Leonardi, for contributions to the data analysis. We also thank the John D. Dingell VAMC and Wayne State University School of Medicine for their ongoing support of this research. Funding of these experiments was made possible by the Department of Veterans Affairs (grant 1I01 RX001095 RR&D Merit Award to AGH); and the National Institute of Health (grant R21DC013895 to MB). The views expressed in this paper do not necessarily reflect the official policies of the Department of Health and Human Services, nor does mention of trade names, commercial practices, or organizations imply endorsement by the U.S. Government.

